# Internal noise measures in coarse and fine motion direction discrimination tasks, and the correlation with autism traits

**DOI:** 10.1101/561548

**Authors:** Edwina Orchard, Steven Dakin, Jeroen J.A. van Boxtel

## Abstract

Motion perception is essential for visual guidance of behaviour and is known to be limited by both internal additive noise (arising from random fluctuations in neural activity), and by motion pooling (global integration of local motion signals across space). People with autism spectrum disorder (ASD) display abnormalities in motion processing, which has been linked to both elevated noise and abnormal pooling. However, to date, the impact of a third limit - induced internal noise (internal noise that scales up with increases is external noise) - has not been investigated in motion perception of any group. Here, we describe a new double-pass motion direction discrimination paradigm that quantifies additive noise, induced noise, and motion pooling. We measure the impact of induced noise on direction discrimination, which we ascribe to fluctuations in decision-related variables. We report that internal noise is higher individuals with high ASD traits only on coarse but not fine motion direction discrimination tasks. However, we report no significant correlations between autism traits, and additive noise, induced noise or motion pooling, in either task. We conclude that internal noise may be higher in individuals with many ASD traits, and that the assessment of induced internal noise is a useful way of exploring decision-related limits on motion perception, irrespective of ASD traits.

## Introduction

Although deficits in social, behavioural and cognitive functioning form the core symptomology of Autism Spectrum Disorder (ASD), sensory and perceptual abnormalities have long been associated with the condition (Asperger, 1944; Grandin, 1992; Kanner, 1943; Kern et al., 2006; O’Neill & Jones, 1997). Sensory issues likely contribute to the complex pattern of behaviours that define ASD, as they are evident in social deficits (facial perception, gestural interpretation, unusual eye contact, difficulties with joint attention) and non-social deficits (light-sensitivity, repetitive/stereotyped behaviours) (Simmons et al., 2009). Differences in sensory processing may play a causative role in core features of autism (Marco, Hinkley, Hill, & Nagarajan, 2011), such as language delay (auditory processing) and difficulty with reading emotion from faces (visual processing). Understanding the mechanisms of such sensory deficits may therefore help to reveal the neural underpinnings of ASD (Zwaigenbaum et al., 2005).

The aetiology of sensory abnormalities in ASD is unknown, but recent work has suggested that higher levels of variability in neural response (internal noise) could be a physiological basis for the condition. Brain imaging studies have shown that individuals with ASD have increased internal noise (Dinstein et al., 2012; Domínguez, Velázquez, & Galán, 2013; Milne, 2011; Weinger, Zemon, Soorya, & Gordon, 2014). Specifically, the variability (but not magnitude) of evoked fMRI response was larger in people with ASD, so that signal-to-noise ratios were lower, across visual, auditory, and somatosensory cortices (Dinstein et al., 2012). Similar differences in neural variability have been reported using resting state magnetoencephalography (MEG; Domínguez et al., 2013), suggesting that high internal noise may represent a fundamental physiological difference in cortical processing of people with ASD (but see, Butler, Molholm, Andrade, & Foxe, 2017; Coskun et al., 2009).

At a behavioural level, the impact of internal noise on visual perception in ASD has mostly been investigated in the motion and orientation domain (Manning, Tibber, Charman, Dakin, & Pellicano, 2015; Manning, Tibber, & Dakin, 2017; Park, Schauder, Zhang, Bennetto, & Tadin, 2017; Zaidel, Goin-Kochel, & Angelaki, 2015). Early evidence for motion processing deficits in ASD indicated that people with ASD were significantly poorer at reporting the perceived direction of stimuli defined by contrast-change than controls, but were normal with stimuli defined by luminance-change (Bertone, Mottron, Jelenic, & Faubert, 2003).

Later research has used motion coherence tasks (Simmons et al., 2009) to measure the minimum number of coherently moving dots (i.e. in a common direction) within a population of randomly moving dots, required to support a reliable report of direction. This work has shown higher motion coherence thresholds in ASD compared to controls (Milne et al., 2002; Spencer & O’Brien, 2006), although not consistently (Brieber et al., 2010; Jones et al., 2011; Manning et al., 2015). Overall, ASD groups exhibit more variable levels of performance compared to controls, speaking to variability within ASD in general (Milne et al., 2006; Pellicano, Gibson, Maybery, Durkin, & Badcock, 2005).-Although behavioural data is equivocal, there is evidence that even when behavioural impairments are absent, neural differences exists between individuals with and without ASD (Brieber et al., 2010; Freitag et al., 2008; Herrington et al., 2007; Kaiser et al., 2010; McKay et al., 2012; Peiker et al., 2015). Could internal noise contribute to such processing differences?

There are two broad explanations for atypical motion coherence thresholds in ASD. First, coherence thresholds could be increased due to poor estimation of local direction, due to high levels of internal noise (Barlow & Tripathy, 1997; Zaidel et al., 2015). Second, coherence thresholds could be increased due to impaired motion pooling (i.e., integration) of local direction signals (Dakin, Mareschal, & Bex, 2005). Pooling local motion signals would combat external (and internal) noise on local motion signals so that a deficit in this process would degrade direction estimation (Dakin et al., 2005; Manning et al., 2015). A recent study investigated these processes in ASD (Manning et al., 2015) and found no evidence for a difference in internal noise between the two groups. However, they did find a difference in motion pooling. Contrary to expectation, individuals with ASD showed more motion pooling when external noise was high, causing ASD children to outperform controls. This finding was, however, not confirmed in a subsequent study, although combining both studies still showed significantly more motion pooling (Manning et al., 2017). These results are consistent with findings (Foss-Feig, Tadin, Schauder, & Cascio, 2013) of enhanced motion perception in ASD compared to controls using a different experimental design.

The equivocal nature of the literature suggests that internal additive noise may not be a strong determinant of motion processing in ASD. However, the ASD literature to date has largely ignored the potential interaction between external noise on internal noise (but see a recent exception, Park et al., 2017, looking at orientation perception). Internal noise can be divided into at least two components: additive and induced (Burgess & Colborne, 1988; Lu & Dosher, 2008). Additive noise is the internal ‘baseline’ level of noise, which is constant across different amounts of input. It is this type of noise that is measured by previous paradigms (i.e., equivalent noise paradigms; Dakin et al., 2005; Lu & Dosher, 2008). Induced noise, on the other hand, is proportional to the amount of noise present in the stimulus (Burgess & Colborne, 1988)^1^. Importantly, when external noise is low (and thus induced noise is low), the main source of internal noise is additive noise. As external noise increases, induced noise increases too, and becomes the main source of internal noise. Figure 1a shows the impact of different types of noise and pooling on thresholds (as measured, e.g., using equivalent noise paradigms). As is clear from this figure, it is difficult to distinguish between the different types of noise. To a large degree (but not completely), the changes in different types of noise are interchangeable, e.g. additive noise and induced noise together may look like a change in motion pooling. It is thus possible that previous studies of motion perception using e.g. equivalent noise paradigms, failed to identify changes in induced noise, because they were interpreted as a change in motion pooling and additive noise.

**Figure 1.**
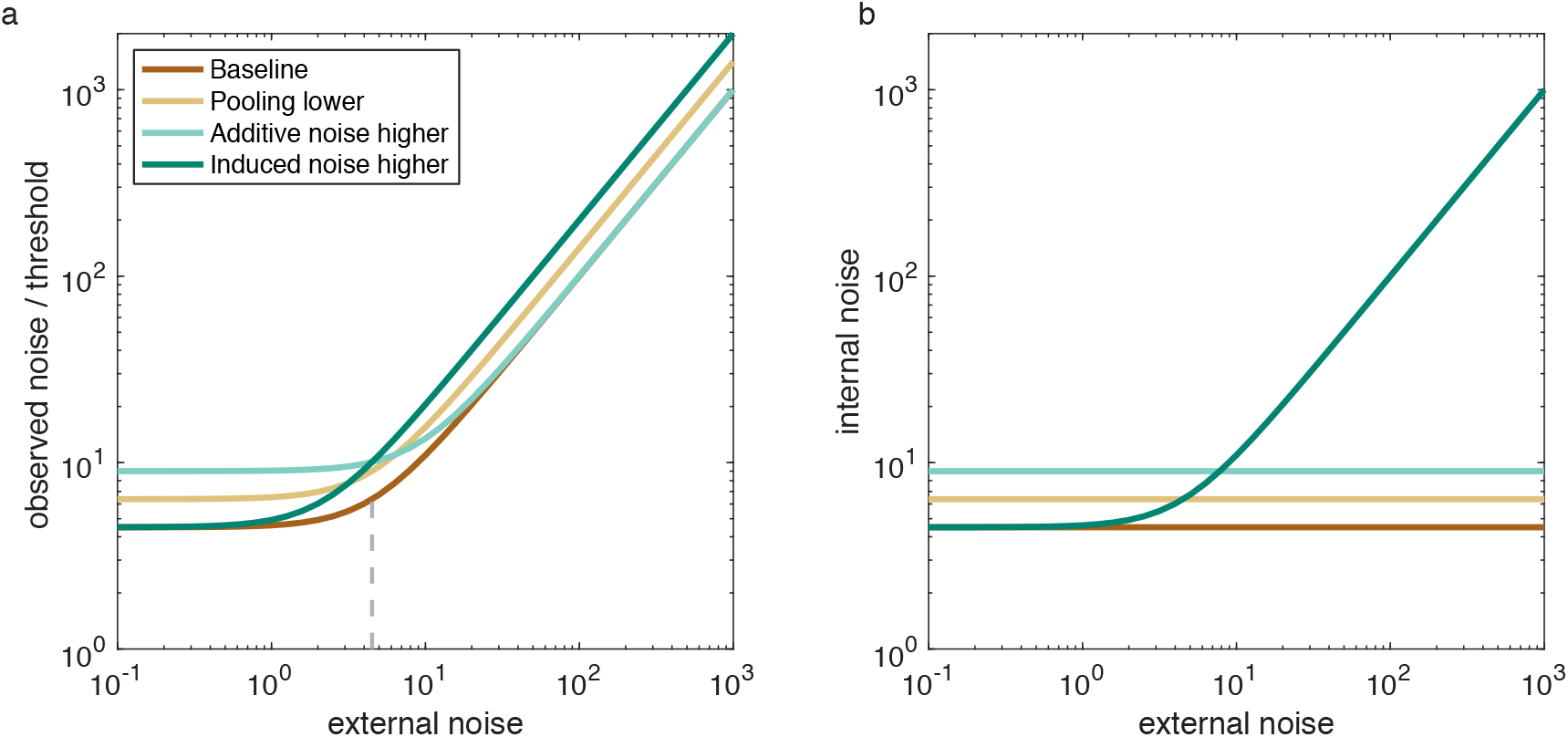
(a) Modelled threshold versus external noise curves, as measured with equivalent noise paradigms. As external (stimulus) noise is increased, performance thresholds (i.e., observed noise) rises. When compared to a baseline (with a certain level of additive, but no induced noise), the following effects are observed: reducing motion pooling leads to a uniform upwards shift in the curve; increasing additive noise increases thresholds but only at low levels of external noise; induced noise increases thresholds especially at high levels of external noise. The vertical dashed line identifies the elbow in the baseline curve, and quantifies the level of additive noise according to the equivalent noise paradigm. (b) Internal noise versus external noise curves, as obtained with a double-pass paradigm. When expressed in terms of the total amount of internal noise, there are clear differences between the different manipulations. Increasing additive noise elevates the internal noise by the same amount independent of the external noise. When induced noise is included, internal noise shows a strong dependence on external noise. Induced noise is the only variable that causes an increase in internal noise in the internal vs external noise plot. In the original approach by Burgess and Colborne (1988) motion pooling was not taken into account. When using this approach, motion pooling scales the curve up or down equally at all external noise levels (causing misestimated noise levels, as shown in this figure). However, when explicitly including motion pooling in the model, motion pooling would have no effect on internal noise estimates (as per our method).

In this study we circumvent this problem by obtaining measures of additive noise, induced noise, and pooling using the double-pass paradigm (Burgess & Colborne, 1988), which is considered to be a direct way of estimating internal noise (Burgess & Colborne, 1988; Lu & Dosher, 2008). In the double-pass paradigm, identical noisy stimuli are presented twice to individuals. A person with high internal noise might perceive such identical stimuli as different (Green, 1964; Haigh, Heeger, Dinstein, Minshew, & Behrmann, 2015) but as the stimuli are identical on both presentations, any perceptual difference must be due to due to internal processes (Green, 1964). The advantage of the double-pass paradigm is that it does not assume that all internal noise is additive and allows one to distinguishes the impact of induced noise from other types of noise (Burgess & Colborne, 1988; Lu & Dosher, 2008) (see Figure 1b). The method has previously been employed to measure internal noise correlations with autism traits in the typically developing population on three non-motion tasks (Vilidaite, Yu, & Baker, 2017), although no estimates of induced noise were derived.

Because the double pass paradigm has not been used for motion discrimination we here further develop it to deal with circular variables. We further tested the influence of additive and induced noise in both a coarse and a fine motion direction discrimination task, because these tasks may depend on different sensory ‘decoding rules’ (Jazayeri & Movshon, 2007), and thus may be differently affected by noise. The difference between fine and coarse motion judgments has not been previously investigated in the context of ASD. However, as ASD is often linked to a more detail-oriented processing, comparing fine and coarse discrimination tasks is potentially very revealing about the underlying mechanisms that are affected in motion processing in ASD.

## Experiment 1. Coarse motion direction discrimination

### Method

#### Participants

Ethics approval was obtained from the Monash University Human Research Ethics Committee (MUHREC), and written informed consent was obtained from all participants prior to participation. This study was completed in accordance with approved guidelines. Participants, 45 healthy adults (31 female, 14 male), ranging from 18 – 40 years old (*M*_age_= 22.07, *SD_age_*= 4.96), were recruited from Monash University Clayton campus. All participants were proficient in English and had normal or corrected-to-normal vision, fulfilling our inclusion criteria. Participants received monetary compensation for their participation. Participants were excluded if the internal noise model (explained below, equation (*1*) fitted with an R^2^ < 0.5. One participant was excluded based on this exclusion criterion.

#### Materials

*The Autism Spectrum Questionnaire* (AQ) was used to measure self-reported autistic traits (Baron-Cohen, Wheelwright, Skinner, Martin, & Clubley, 2001). The AQ comprises 50 items on a four-point Likert scale (*definitely agree, slightly agree, slightly disagree, definitely disagree*) and was administered and scored on a computer.

We also collected data from the *Kaufman Brief Intelligence Test second edition* (KBIT-2; Kaufman & Kaufman, 2004). Because our cohort included a large proportion of non-native English speakers, this test did not provide accurate estimates of the intelligence quotient, but all individuals scored > 70.

#### Apparatus

This study was completed in an experimental room without external lights, with artificial lights turned on. Participants sat comfortably in front of the computer, with their head stabilised by a chin rest. The experiment was displayed on a VIEWPixx/3D monitor, with a refresh rate of 120Hz and a resolution of 1920 x 1200 pixels, and viewed from a distance of 114 cm. All experimental displays (including stimuli and AQ administration and scoring) were created using Matlab and OpenGL, with PsychToolbox extensions (Brainard, 1997; Pelli, 1997).

#### Stimuli

We used random-dot motion stimuli (Figure 2), made up of 200 circular dots (100 black, 100 white, diameter 0.07°) moving within a circular aperture, which was outlined in black against a grey background. A red fixation mark was provided (diameter 0.35°), which was surrounded by a 0.67° exclusion zone, in which no dots were drawn in order to decrease potential pursuit eye-movements. Each individual dot moved (speed = 2.38°/sec, lifetime = 8 frames) in a direction randomly chosen from a wrapped normal Gaussian distribution (mean 22° clockwise or anticlockwise from vertical, with standard deviations of: 0, 35, 45, 60, 70, 80, 90 or 100 degrees. A noise level of zero means that all dots move in the same direction (22° clockwise or anticlockwise of upwards motion).

**Figure 2.**
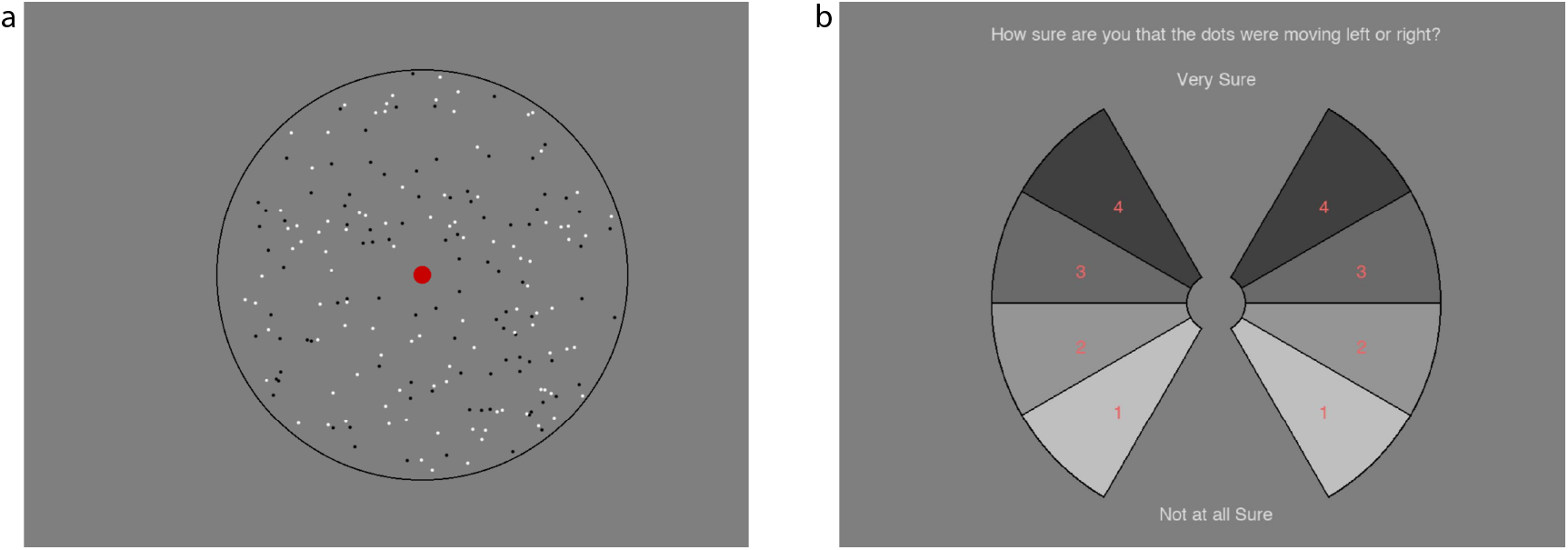
Screenshot of visual motion task stimuli (a) and response screen (b). Participants viewed the stimuli and then gave a response, clicked on a wedge in the response screen, indicating whether the dots are moving to clockwise or anti-clockwise of vertical, and how confident they were in their response.

Participants were required to indicate the perceived direction of the motion stimuli using a response screen (Figure 2b). This consisted of a confidence ‘wheel’, comprising eight numbered wedges. The four wedges on the left or right, were used to indicate that the dots were, on average, perceived to move anti-clockwise or clockwise of vertical, respectively. Each of the wedges was numbered (1= *Not at all sure*, 4= *Very Sure*) and shaded (1=lightest, 4=darkest) and the confidence screen was flipped up-side down every 100 trials.

#### Procedure

Participants performed the KBIT-2, which took approximately 40 minutes, then the visual motion task, and finally the AQ questionnaire.

In the motion task, stimuli (Figure 2a) were presented for 0.75 seconds followed by the response screen (Figure 2b). Participants were instructed to focus their gaze on the fixation point (red dot) at the centre of the screen during stimulus presentation and were asked to judge the average direction of the moving dots. Participants indicated their decision by a clicking on a wedge in the response screen which served to indicate if they thought dots were moving clockwise or anti-clockwise of vertical, and how confident they were in this judgment (Figure 2b).

Our study employed a double-pass paradigm, similar to that used by Burgess and Colborne (1988), a method whereby two identical presentations (passes) of each stimulus is made over two separate trials. There were 100 unique stimuli, and therefore 200 trials in total, for each noise level. The first passes were run first in random order, and the second pass were then run in the same order. Participants were given self-timed breaks after every 100 trials. A total of 1600 trials were run.

### Data Analysis

Analyses were run in MATLAB (Mathworks Ltd) and Bayesian statistics were performed using the jsq module from the JASP team run in Jamovi (jamovi, 2020; JASP Team, 2018).

We employed a double-pass paradigm to quantify internal noise (Burgess & Colborne, 1988). Values for internal additive and induced noise and motion pooling were calculated based on participants’ performance on the visual motion task. Internal noise can be estimated by examining the accuracy (over individual trials) and consistency (between the two passes) of a person’s response (Burgess & Colborne, 1988). For this task, a person is accurate if the dots are moving left (or right) and the participant responds “left” (or “right”). A person is consistent if they choose the same direction on both test and retest presentations of a trial, regardless of whether or not they were correct (and regardless of their reported confidence).

#### Internal noise

When noise levels are expressed as a ratio of internal over external noise (σ_int_/σ_ext_), there exists a fixed (non-linear) relationship between accuracy and consistency at different levels of σ_int_/σ_ext_ (Burgess & Colborne, 1988). These relationships are shown in Figure 3 for different ratios of σint/σext (grey dashed lines). Internal noise values can be calculated by fitting such a curve to experimental data, and finding the ratio σint/σext that best captures the data. Then, because σext is known, σint can be calculated. Because this approach will not work for circular variables, we calculated the curves numerically for wrapped circular distributions. Figure 3 shows a comparison of the non-circular (coloured lines) and circular approach (gray dashed lines), using identical parameters. When noise is large (towards the lower side in the plot), circular data deviate from the non-circular data. An unfortunate consequence of the circular nature of the data, is that the exact shape of the curves is dependent on the signal strength. When the mean angle (i.e., signal) is small, the data are well approximated by an approach using Gaussian distributions (only showing deviations at very large noise values).

**Figure 3.**
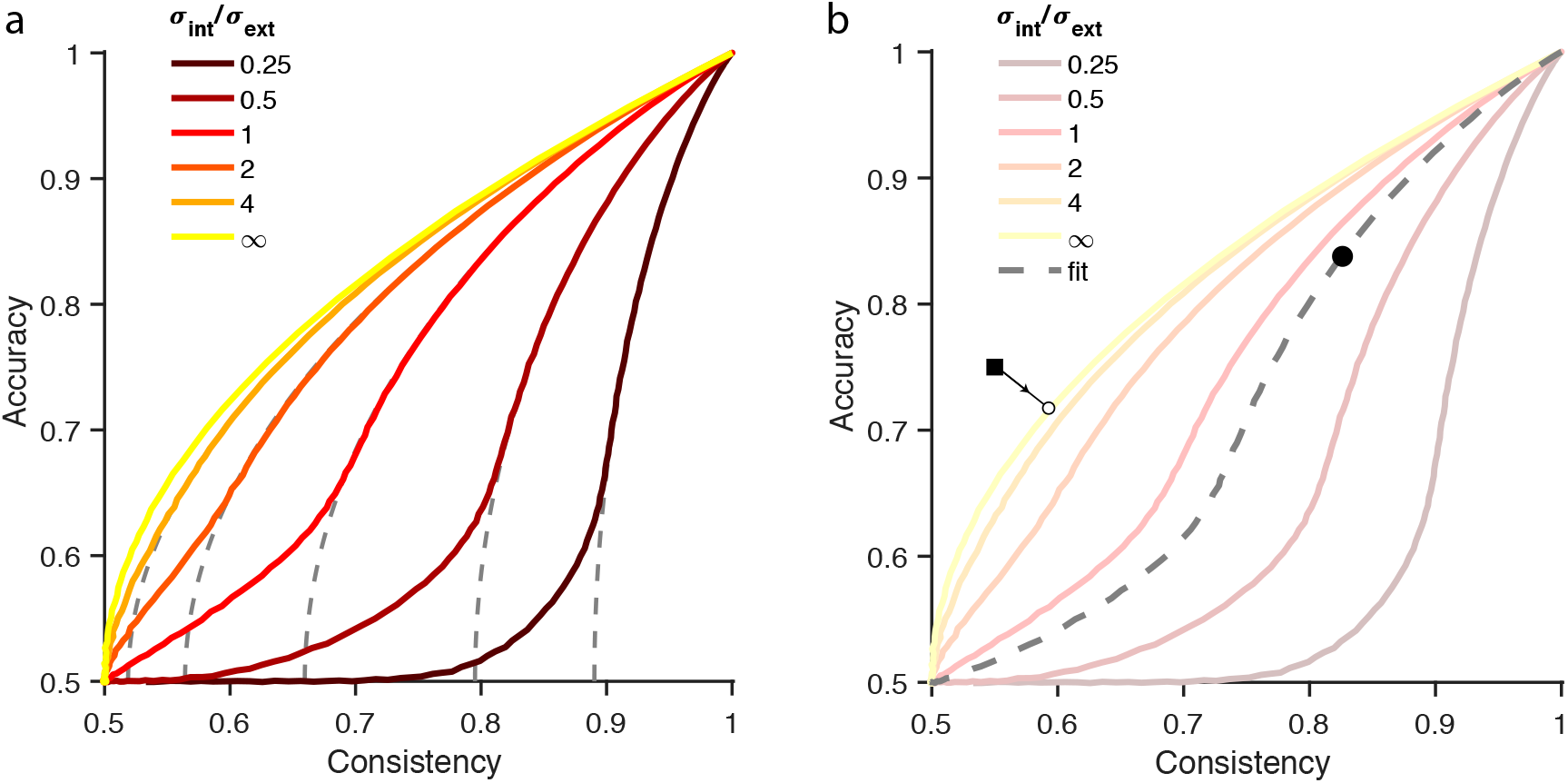
Relationship between accuracy and consistency, dependent on different ratios of σ_int_/σ_ext_. (a) Coloured lines are calculated using a wrapped Gaussian distribution, dashed grey lines are based on Gaussian distributions used by Burgess and Colborne (1988). The line for σ_int_/σ_ext_ = ∞ is the case without external noise. (b) Schematic of how data is analysed for two example data points. Data falling above the σ_int_/σ_ext_ = ∞ curve, is mapped onto the nearest point on the σ_int_/σ_ext_ = ∞ curve, and the internal noise associated with that point is taken as the measured internal noise. For all other data (circle) the best fitting line determines σ_int_/σ_ext_.

Instead of fitting the data, we constructed a large lookup table for accuracy-vs-consistency curves ratios σ_int_/σ_ext_ of (1.12^q^-1)/25, with q ranging from 1 to 59 in steps of 0.5. Individual data points were compared to these curves, and the curve to which the data point showed the smallest squared Euclidian distance was taken as the best fit (Figure 3b). The squared distance was calculated taking into account deviations in terms of both accuracy and consistency. When the participant’s data showed an accuracy <0.5 it was adjusted to (1-accuracy), so that fits could be made. This happened 6 times (out of 45 × 8 = 360 data points, i.e., 1.67% of all data points). This procedure determined σ_int_/σ_ext_, as well as σ_int_ (since σ_ext_ is known). When a data point fell above the curve σ_int_/σ_ext_ = ∞ (e.g., square in Figure 3b), internal noise was estimated by finding the point on the internal-noise-only curve that was closest to the data point (square in Figure 3b, which was mapped to the open circle). The internal noise value that was associated with that point on the curve was taken as the internal noise estimate for our data point. Simulations showed that this resulted in correct approximations of internal noise levels (see *Simulations* in https://osf.io/4gdkt/).

#### Determining Additive and Induced internal noise

In equivalent noise paradigms, the amount of internal (additive) noise is determined by finding the elbow in the curve in Figure 1a, dashed line. This approach however, ignores the possible contributions of induced noise. Therefore, if induced noise influences the task at hand, estimates of additive noise can be incorrect. For example, when introducing induced noise in the data of Figure 1a, the elbow moves to lower external noise values, thus underestimating the amount of additive noise. Unfortunately, it is difficult to determine from the plots in Figure 1a, whether induced noise is present, because the curves are very similar (and can be made nearly overlapping by appropriately setting additive noise and pooling parameters). However, using the double-pass paradigm, one can plot the internal noise versus the external noise values (Figure 1b). Induced noise can be derived from the increase in internal noise that depends on the level of external noise, with a slope of 0 indicating that there is no significant influence of induced noise. This makes intuitive sense, as the internal noise only consists of a fixed level of constant additive noise, which is not changed by the level of external noise, thus tracing a flat line in the internal versus external noise plot.

According to Burgess and Colborne (1988), the induced noise σ_ind_ is directly proportional to the external noise, and thus σ_ind_ = *m* σ_ext_. The total internal noise is σ_int_ = sqrt(σ_add_^2^ + σ_ind_^2^) = sqrt(σ_add_^2^ + (*m* σ_ext_)^2^).

The induced noise factor *m* can be derived in various ways, and because we had no *a priori* reason to assume which one worked best, we performed a simulation study (see Appendix 1). This simulation study showed that the best way to determine additive and induced noise was to fit the following function to the collective internal noise data per participant:

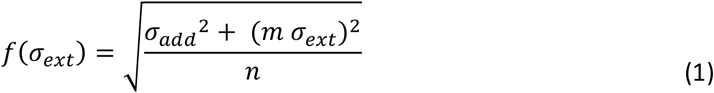

Where n is the number of motion samples that are taken to estimate motion direction, i.e. motion pooling. We fitted equation (1 to individual participant data, after first independently estimating n from the data (see below).

#### Determining motion pooling

Better performance in a task may not only result from lower noise levels, but also when the direction information from multiple dots is combined. This is called (global) motion pooling (Dakin et al., 2005). To derive motion pooling in our task, we plotted iso-external noise lines (Figure 4). These lines trace, for one value of external noise, the expected values of consistency and accuracy. Note that these lines are dependent on the parameters of our stimulus, and thus will be different in different experiments, like the fine discrimination task below. Each data point will fall on only one curve, and this curve indicates the *observed* external noise. The data generally fall on a curve that has a lower level of observed external noise than the amount of external noise that was actually present in the stimulus, indicating that information from multiple dots is pooled (see e.g., the data points from σ_ext_ = 80, which falls on the line of observed external noise = 30; Figure 4).

**Figure 4.**
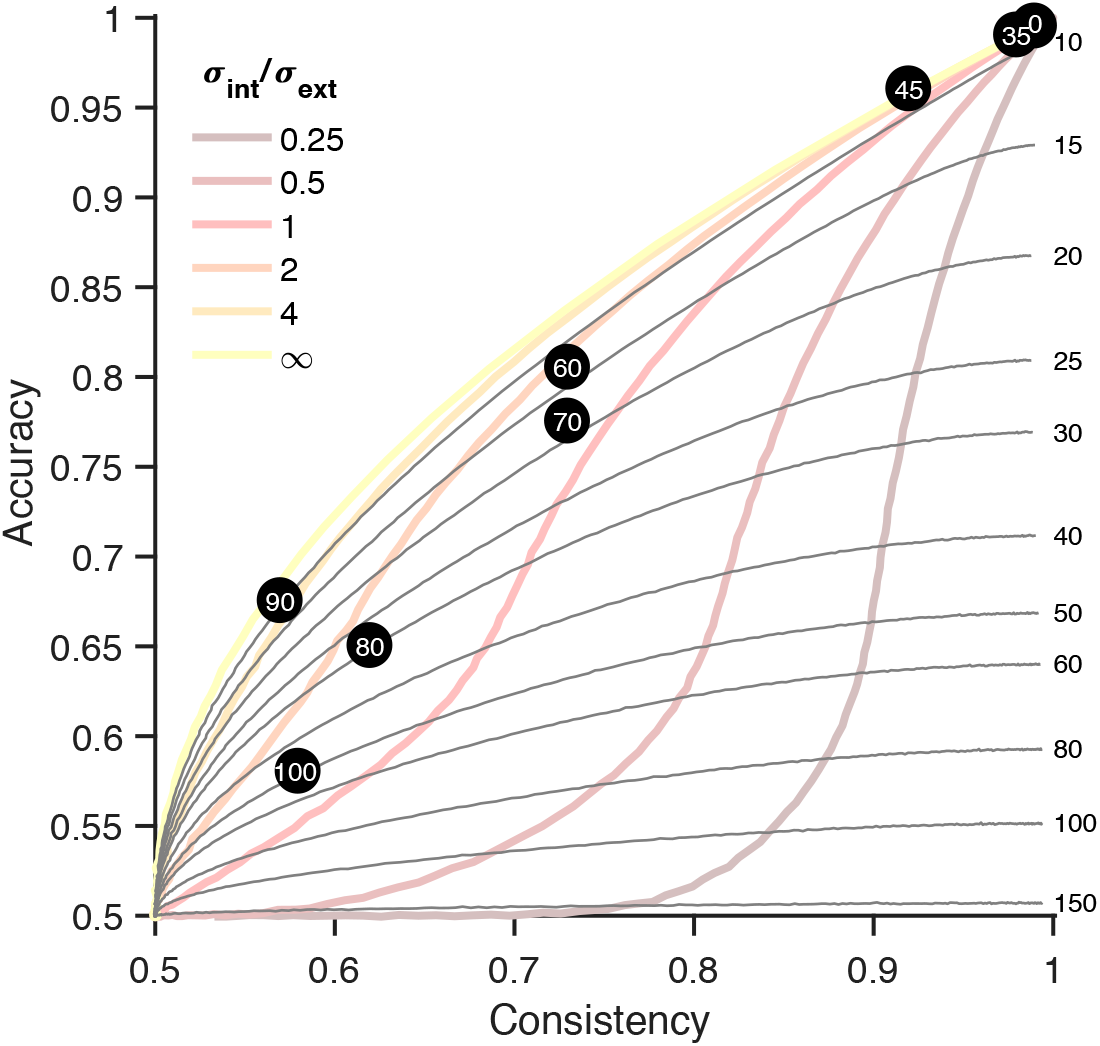
Observed external noise and motion pooling. This plot has the same layout as Figure 3, but also shows iso-external noise lines: the accuracy and consistency expected for a given external noise level (displayed on the right). The black dots are experimental data for one participant, with the external noise for each condition indicated within the dot. The external noise for each condition is higher than the iso-external noise line it falls on, suggesting that the observed/effective external noise is lower than the actual external noise, an indication that motion pooling was occurring.

The level of motion pooling is derived with a characteristic of the central limit theorem that:

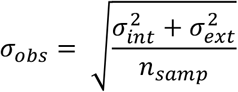

where σ_obs_ is the observed standard deviation of the response, and *n_samp_* is the effective number of samples that are combined to give a motion direction estimate (i.e. motion pooling). In our case:

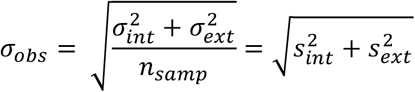

Where σ_int_ and σ_ext_ are the real internal and external noise values, and s_int_ and s_ext_ are observed values. The observed values have the motion sampling already taken into account, which is why *n*_samp_ does not appear on the right-hand side of the equation. Since we know the ratio (a) between internal and external noise and thus σ_int_ = α * σ_ext_, this equation can be rewritten as

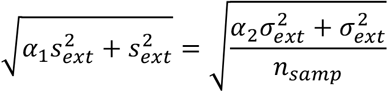

and

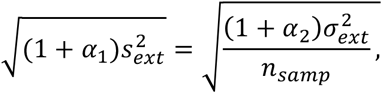

which can be rewritten as:

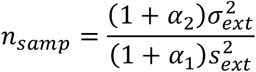

Assuming that α_1_ = α_2_, i.e. that the motion pooling does not affect the ratio of internal to external noise, we can derive motion pooling as:

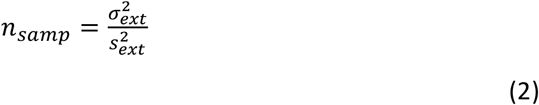

Since we know both σ_ext_ (the external noise, i.e., standard deviation of directional noise) and sext (the observed external noise), we can calculate the motion pooling *n*_samp_. Simulations showed that this method of estimation works, but only for the lower noise levels (we used 35, 45, 60 degrees) for our coarse discrimination task, and for the higher noise levels for the fine discrimination task (we used 3.71 – 51.19). These simulations can be found in *Simulations* at https://osf.io/4gdkt/.

For data points that lay beyond the σ_int_/σ_ext_ = ∞ curve, observed external noise could not be determined, and these data points were consequently not used to estimate motion pooling.

### Results

#### Influence of external noise on accuracy, consistency and confidence

As external noise increased, accuracy, consistency and confidence ratings decreased, reflecting increased task difficulty (Figure 5). In particular, both accuracy and consistency decreased from near-perfect to near-chance levels. This suggests that the spread of noise levels measured both the upper and lower limits of participant performance. Although we do not present any further analysis of confidence ratings here, confidence data have been deposited in a large openly available confidence database (Rahnev et al., 2020).

**Figure 5.**
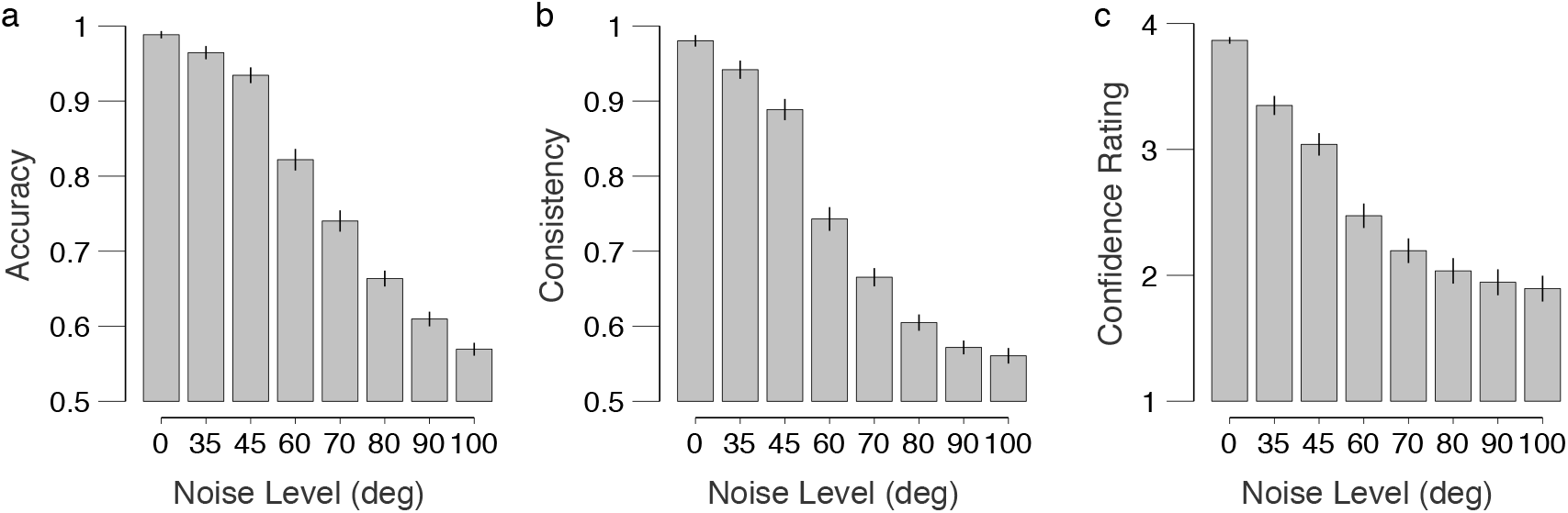
Mean behavioural performance. (a) Accuracy, (b) consistency, and (c) confidence decrease as external noise level increases, but remain significantly above chance level at all noise levels. Error bars are standard error of the mean calculated over participants.

We investigated whether AQ had a significant effect on the dependence of accuracy and consistency on noise. Numerically, the accuracy and consistency were lower in people with higher AQ scores. To test the statistical significance of this finding, we constructed an unrestricted linear model with AQ, noise level and their interaction as terms, and we compared them to models without the interaction, and models without both the interaction and the AQ term. Likelihood ratio tests revealed no significant differences between unrestricted models and those excluding the interaction (accuracy: χ^2^(1) = 0.07, *p* = .78; consisteny; χ^2^(1) = 0.68, *p* = .41), and between unrestricted models and those exluding both AQ and the interaction (accuracy: χ^2^(2) = 3.51, *p* = 0.17; consistency: χ^2^(2) = 4.08, *p* = .13).

However, comparing the models with AQ (but without interaction) and those with only external noise, showed near significant results (accuracy: χ^2^(1) = 3.44, *p* = .065; consistency: χ^2^(1) = 3.40, *p* = .065), suggesting that AQ may have a small influence, but our current experiment was not powerful enough to reveal it. Overall, there appears to be no statistically significant influence of AQ on the accuracy, or consistency of participants’ report.

#### Internal noise

Internal noise was derived from the accuracy-consistency plots (see *Methods*) and depended on the external noise (Figure 6). The plot shows that internal noise values depend on external noise, a sign of the involvement of induced noise. A repeated-measures one-way ANOVA with external noise as the factor was significant (internal noise transformed as log(σint+1), *F*(2.28, 98.21) = 220.68, *p* < 0.0001, η_p_^2^ = 0.84, Greenhouse-Geisser corrected). When performing median split on the AQ scores (median = 18), we obtain a group of *n* = 20 that scored AQ < 18, and a group of *n* = 20 that scored AQ > 18 (we discard the subjects with a median score for this analysis). The mixed-design one-way repeated-measured ANOVA with the factor external noise, and group (Low vs High AQ) showed a significant effect of external noise (*F*(2.22, 84.49) = 205, *p* < .0001, *η*_p_^2^ = 0.84, Greenhouse-Geisser corrected), and AQ group (*F*(1, 266) = 4.33, *p* = .044, *η*_p_^2^ = 0.10), with internal noise increasing as external noise increased, and higher internal noise for the high AQ group. The interaction was not significant (*F*(7,266) = 0.21, *p* = 0.98).

**Figure 6.**
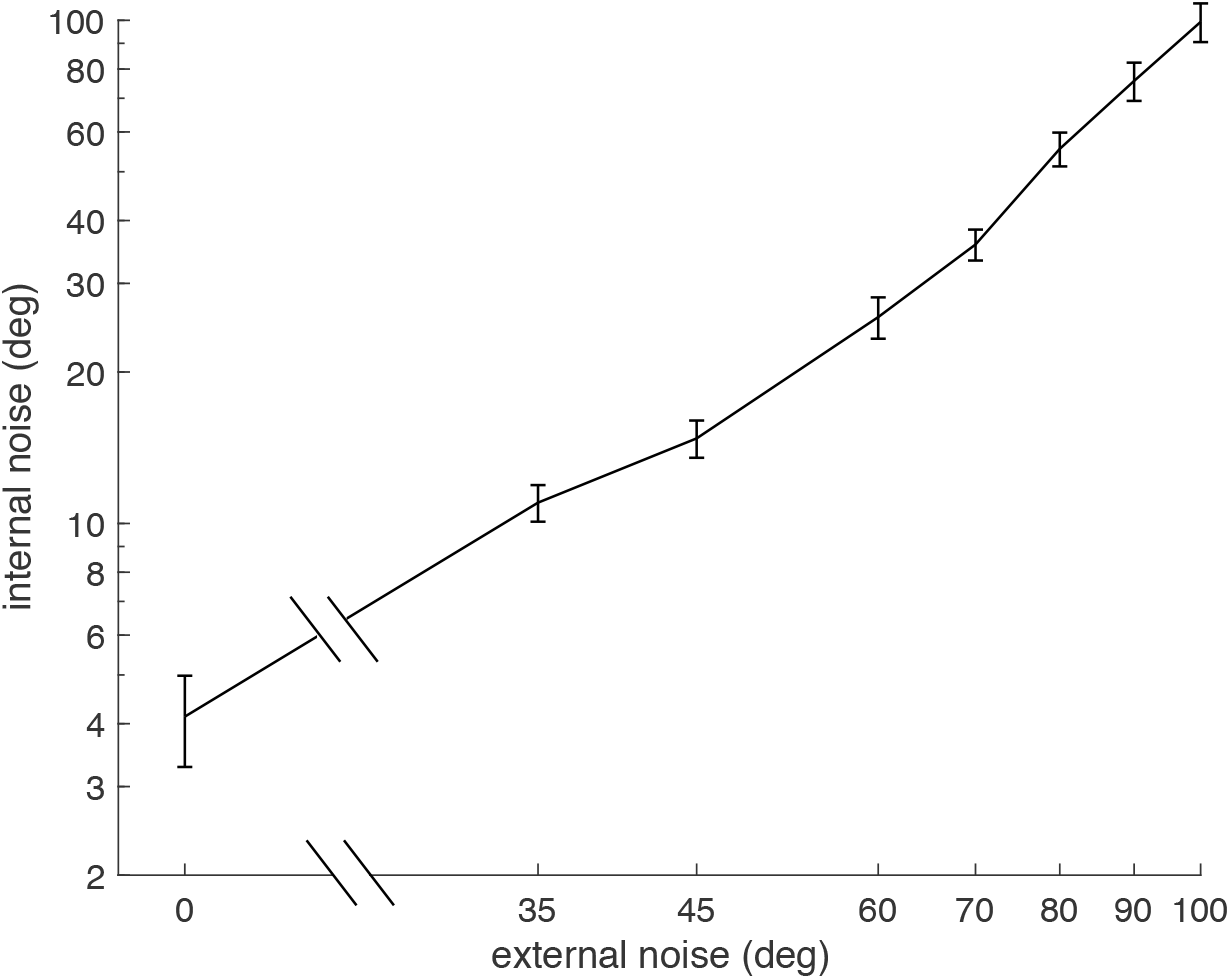
Dependence of estimated internal noise on physical external noise. Values are averaged across participants, and error bars show indicate ± s.e.m.

#### Additive noise

We estimated additive and induced noise by fitting Equation (*1* to individual participant data. Unfortunately, our design did not support accurate estimation of additive noise, because the task was too easy, resulting in (near) perfect performance. Our simulations indicated that internal additive noise could not accurately be estimated from this experiment, so we did not perform statistical analyses on it. However, the individual data are shown in Figure 7a.

**Figure 7.**
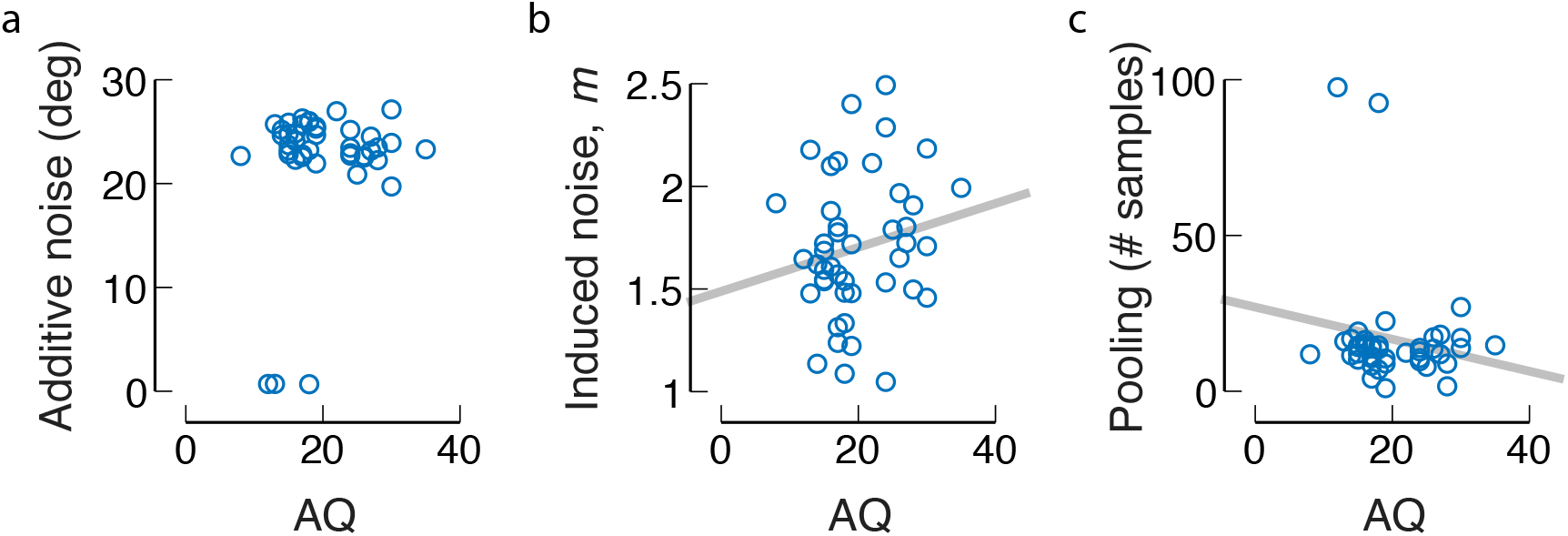
Correlations for the coarse discrimination task. (a) additive internal noise, (b) the induced noise factor m, (c) motion pooling, dependent on AQ. One outlier was removed in (c).

#### Induced noise

Internal noise estimates increase with increasing external noise (Figure 6), which is a sign of the involvement of induced noise. To obtain insight into whether this involvement differs between individuals with different levels of AQ, we estimated individual levels of induced noise by fitting equation (*1*. The average induced noise factor over participants was 1.70 (median 1.66), which is significantly larger than 0 (one sample t-test, *t*(43) = 33.12,*p* < 0.0001, Cohen’s *d* = 4.99). These values did not correlate with the AQ measure (Kendall’s tau-b = 0.09, *p* = 0.39; Bayesian correlation Kendall’s tau 0.094, BF_10_ = 0.29).

#### Motion pooling

We computed the motion pooling at each level of external noise for each participant, using equation (*2*. We then took the median value over the calculated pooling values from external noise conditions 35, 45, 60 degrees, discarding conditions in which pooling could not be estimated. These particular external noise values were chosen based on simulations (see methods, and Appendix 1). The median value was 13.73 samples, which was significantly higher than 1 (Wilcoxon signed rank test, p < .0001) indicating that participants were responding using direction information from multiple dots. We also note that 13.73 is rather close to the square root of the number of samples present 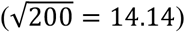, which has previously been proposed as a simple rule of thumb for estimating effective sample eize in averaging tasks (Dakin, 2001). There was no correlation between the extent of motion pooling and AQ (Kendall’s *tau* = −0.08, *p* = 0.45, Figure 7c; one multivariate outlier was removed based on Mahalanobis distance > 13.8155). A Bayesian correlation suggested moderate evidence for the null hypothesis (Kendall’s *tau* = −0.083, BF_10_ = 0.27).

## Experiment 2. Fine discrimination task

### Methods

#### Participants

We recruited 37 healthy adults from Monash University Clayton campus. All participants were proficient in English and had normal or corrected-to-normal vision, fulfilling our inclusion criteria. Participants received a monetary compensation for their participation. Participants were excluded if the internal noise model (equation (*1*) fitted with an R^2^ < 0.5. Ten participants were excluded on this basis, leaving 27 participants in the final sample (20 females, 7 males, ranging from 18 – 32 years old, *M*_age_ = 24.8, *SD*_age_= 5.33).

#### Stimuli

Stimuli and methods were identical to the coarse discrimination task except that the mean motion direction was ±5 degrees from vertical, and the standard deviations of the Gaussian direction distributions were 0, 2.2, 3.7, 6.3, 10.6, 17.9, 30.3, and 51.2 degrees.

#### Analyses

All analyses were as in experiment 1, except that lookup tables for analyses were recalculated for a signal value of 5 degrees from vertical.

### Results

#### Influence of external noise on accuracy, consistency and confidence

As in the coarse discrimination task, the fine discrimination task showed decreases in accuracy, consistency and confidence rating as external noise increased, reflecting increased task difficulty (Figure 8). Numerically, the accuracy and consistency were lower in people with higher AQ scores. To test the statistical significance of this finding, we constructed an unrestricted linear model with AQ, noise level and their interaction as terms, and we compared them to models without the interaction, and models without both the interaction and the AQ term. Likelihood ratio tests revealed no significant differences between unrestricted models (incuding AQ, noise level and their interaction as terms) and those excluding the interaction (accuracy: χ^2^(1) = 0.068, *p* = .79; consisteny; χ^2^(1) = 0.13, *p* = .72), and unrestricted models and those exluding AQ and the interaction (accuracy: χ^2^(2) = 0.07, *p* = .96; consistency: χ^2^(2) = 1.15, *p* = .56). Comparing the models with AQ (but without the interaction) to those with only external noise showed no significant effects either (accuracy: χ^2^(1) = 0.007, *p* = 0.93; consistency: χ^2^(1) = 1.02, *p* = .31). Overall, there appears to be no influence of AQ on the accuracy, and consistency in our fine motion discrimination task.

**Figure 8.**
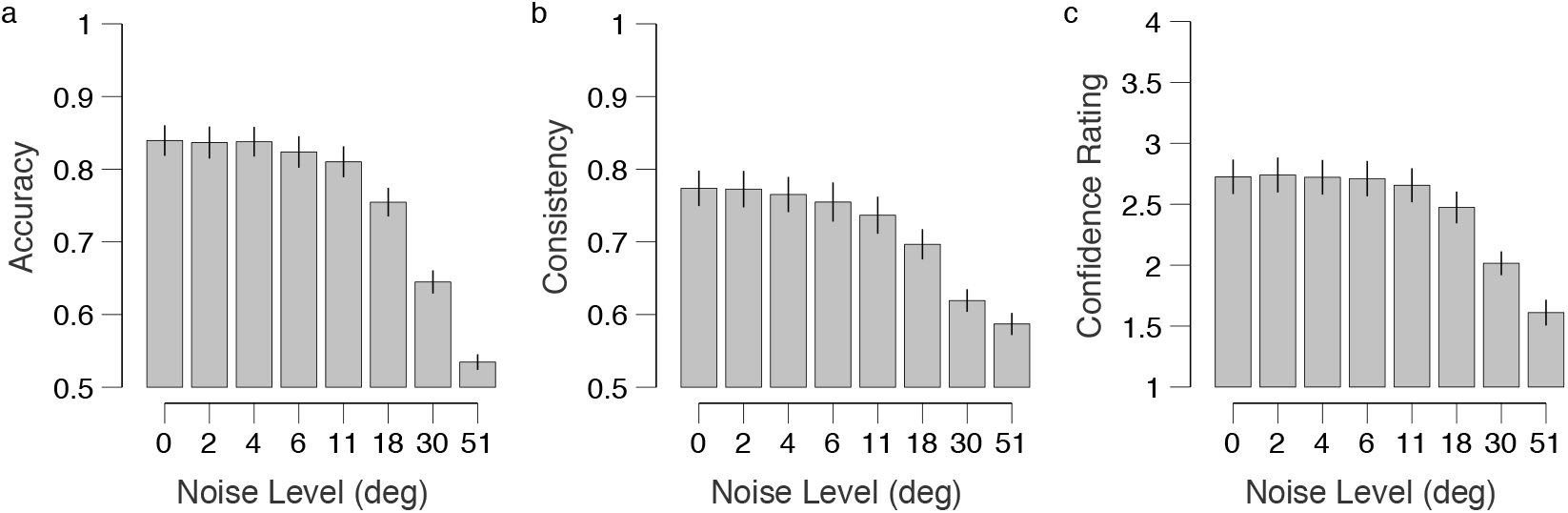
Mean behavioural performance in the fine discrimination task. (a) accuracy, (b) consistency, and (c) confidence, all decrease as external noise increases, but remain significantly above chance at all noise levels. Error bars show the standard error of the mean over participants.

#### Additive noise

In contrast to Experiment 1, we were able to obtain individual additive noise estimates from participants performance of the fine discrimination task. Additive noise, taken as the observed internal noise when external noise was 0, was larger than 0 (Mean ± SEM of sample: 15.24 ± 1.27, median = 14.17, *t*(26) = 11.97, *p* < .0001, Cohen’s *d* = 2.30). There was no significant correlation between our individual additive noise estimates and corresponding AQ scores (Kendall’s *tau* = −0.05, *p* = 0.72; BF_10_ = 0.25; Figure 10a).

**Figure 9.**
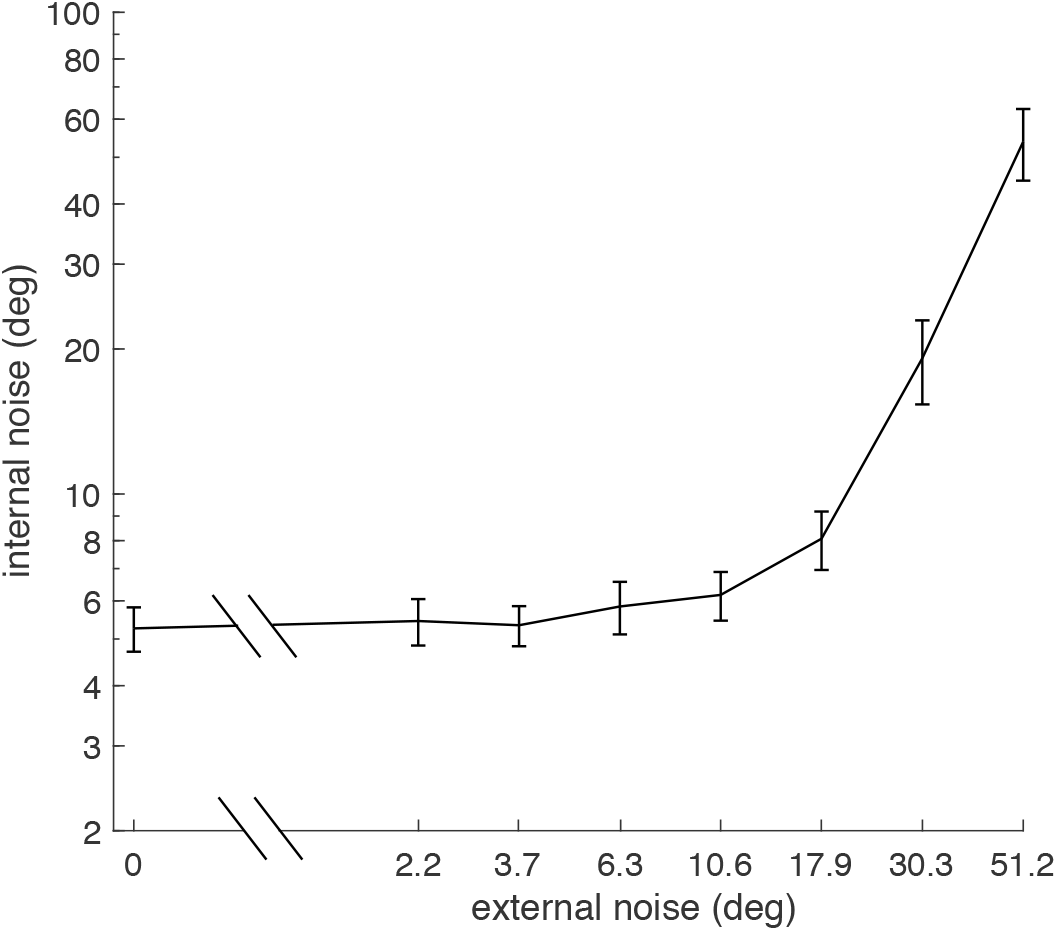
Dependence of mean (± s.e.m.) internal noise on external noise for the fine discrimination task, averaged over participants. Datapoints which were estimated as 0 internal noise were ignored.

**Figure 10.**
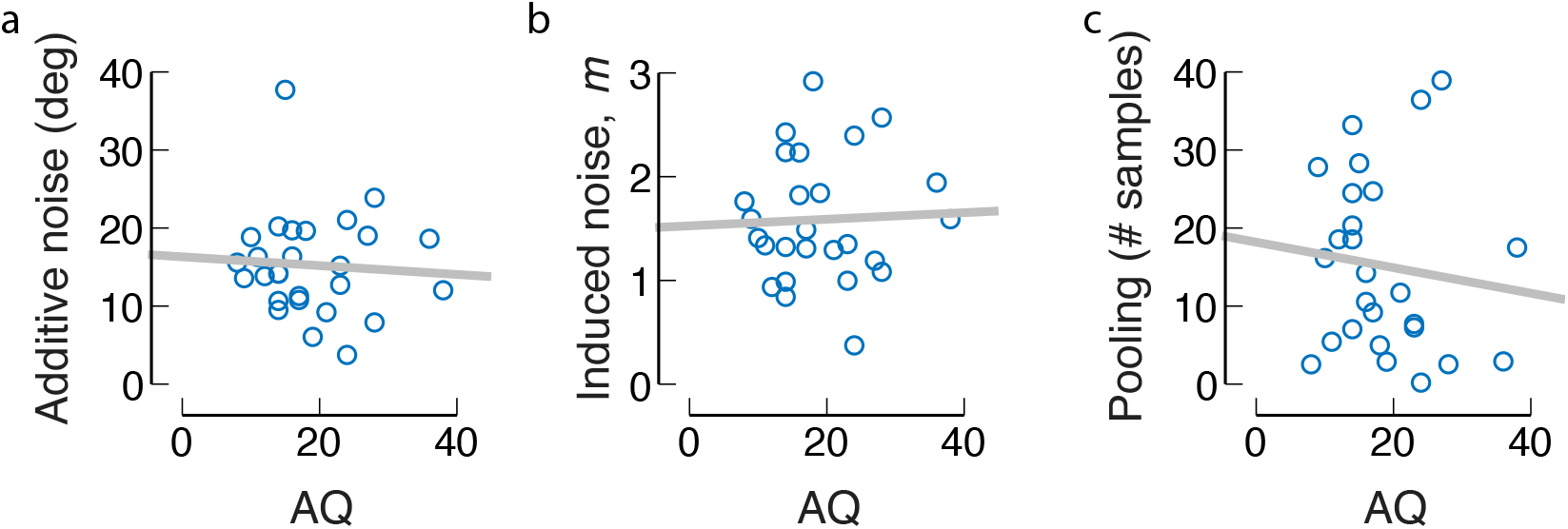
Estimates of (a) additive internal noise, (b) the induced noise factor m, (c) motion pooling, from the fine direction estimation task, plot against AQ.

#### Induced noise

The influence of induced noise was again shown by an increase in internal noise with increasing external noise (Figure 9). Mean induced noise factor, *m*, was 1.72 (median 1.49), which is significantly larger than 0 (one sample t-test, *t*(26) = 9.76, *p* < .0001, Cohen’s *d* = 1.88). These values were not correlated with the AQ measure (Kendall’s *tau-b* = 0, *p* = 1; BF_10_ = 0.25; Figure 10b; one multivariate outlier was removed).

#### Motion pooling

Mean pooling was 18.56 samples (median 14.24), which was significantly higher than 1 (Wilcoxon signed rank test, p < .0001). There was no correlation with AQ (Kendall’s *tau* = - 0.13, *p* = 0.36; Figure 10c; one multivariate outlier was removed). A Bayesian correlation suggested moderate evidence for the absence of a correlation (BF_10_ = 0.28).

#### Comparing fine and coarse discrimination tasks

From Figure 6 and Figure 9, it appears that the internal noise is larger in the fine discrimination task than in the coarse discrimination task. We compare internal noise between the two tasks at similar external noise conditions. We found that internal noise is significantly larger in two of the three comparisons (Wilcoxon rank sum test; at σ_ext_ = 0, *Z* = 2.24, *p* = .025; at fine σ_ext_ = 30.5, coarse σ_ext_ = 35, *Z* = 1.10, *p* = .27; at fine σ_ext_ = 51.2, coarse σ_ext_ = 45, *Z* = 5.80,*p* < .0001). Mean induced noise was not different between the two tasks (*Z* = 1.00,*p* = .32, Wilcoxon rank sum test). Comparing the motion pooling data between coarse and fine discrimination tasks showed no significant difference (*Z* = 0.05, *p* = .96, Wilcoxon rank sum test).

## Discussion

We investigated how fine and coarse motion discrimination is limited by internal noise (both additive and induced), and by pooling (or multiplicative noise), and we also examined if these limits were correlated with Autism Spectrum Disorder (ASD) traits in a typically developed adult population. We found evidence for higher internal noise in ASD when performing a coarse discrimination task, but not in a fine discrimination task. However, in neither the coarse and fine discrimination task were additive noise, induced noise, or motion pooling correlated with ASD traits.

### Additive noise influences on motion perception

Several studies have reported high internal noise in ASD populations compared to control groups using brain-imaging techniques such as functional magnetic resonance imaging (fMRI; Dinstein et al., 2012; Haigh et al., 2015), electroencephalography (EEG; Milne, 2011; Weinger et al., 2014) and magnetoencephalography (MEG; Ishikawa, Shimegi, & Sato; Peiker et al., 2015). These results led us to anticipate a positive correlation between AQ score and additive noise which we did not observe in the fine discrimination task, and which we were unable to estimate in the coarse discrimination task. Additive noise has not been previously investigated across AQ in a typically developed population, and our results suggest that additive noise does not vary with traits of ASD in the general population. These results are consistent with other research that did not find significant differences in additive noise between ASD and control groups, in both behavioural (Manning et al., 2015) and brain-imaging (Coskun et al., 2009) studies.

Aside from the lack of a link to autism traits, our results do speak to the involvement of additive noise in the perception of motion. We found considerable differences in the amount of additive internal noise between individuals. These individual differences in noise were correlated with (and in fact, calculated from) differences in performance measures (accuracy and consistency) in a motion discrimination task. This suggest that internal additive noise can determine task performance differences between individuals, although at the moment it appears that it cannot stratify individuals along the broader ASD spectrum.

### Induced Noise

We identified a strong influence of induced noise on motion direction discrimination. In motion tasks, induced noise has not been previously investigated in relation to ASD, either between ASD and control groups, or across AQ score within the broader spectrum. Instead, studies have investigated internal noise without splitting it into additive and induced noise. For example, Manning et al. (2015) varied the amount of external noise in a motion discrimination task but only looked at two levels of noise. Overall, they found better performance at high noise levels for individuals with ASD relative to the control group, but no difference at low noise levels. The authors tentatively attributed performance differences at high noise levels to differences in motion pooling (Manning et al., 2015), but because motion pooling should also affect performance at low noise levels, this explanation was speculative. However, differences in induced noise provide an alternative explanation for this difference. For example, lower induced noise yields better performance at high external noise levels but not at low external noise levels (see e.g. Figure 1b), which is consistent with experimental findings in ASD (Manning et al., 2015; Manning et al., 2017). Our own results do not directly support this interpretation, however, as we did not find a correlation between induced noise and AQ. Perhaps we lacked enough individuals with high AQ scores to show a clear dependence. Alternatively, large differences in induced noise may only appear when comparing a control group to a group with a clinical diagnosis of ASD.

Burgess and Colborne (1988) concluded, using a model introduced by Wilcox (1968), that induced noise could be explained by fluctuations in a decision-variable, if the standard deviation of the fluctuations (i.e., noise) in the decision variable is proportional to the signal to noise ratio (and thus to external noise, when *σ_ext_ ≫ σ_int_*). This relationship can be derived as follows: variability in the decision variable (σdec) adds to the variability in the response (and thus increases threshold), just like additive noise does. Thus, total noise would be 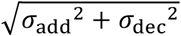. Now, if we assume that σ_dec_ is proportional to the external noise, then total noise is 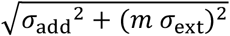, which is the definition of induced noise.

This analysis suggests that induced noise could reflect fluctuations in a decision variable, e.g. the criterion, or an internal standard. It rests on the assumption that decision variable is proportional to the signal to noise ratio, but is that warranted? Past research has shown that decision/criterion noise increases when, within an experiment over trials, a wider range of stimulus values is tested (Gravetter & Lockhead, 1973). We reason that a wider stimulus range over trials may have similar effects to a wider stimulus range within a trial (that is, stimulus noise), which implies that more stimulus noise would translate into more decision/criterion noise. This is supported by research that shows that more noisy stimuli lead to less decision confidence (which is probably related to criterion noise), even when performance (in terms of d’) is equated (Spence, Dux, & Arnold, 2016).

Past research in non-motion tasks (Vilidaite et al., 2017) is consistent with interpretation that induced noise may be linked to decision noise. It was found that internal noise measured through the double-pass paradigm correlated strongly in three quite different tasks (contrast, and face discrimintation, and a mathematical task), and the priniciple component across the internal noise measures from these three tasks was correlated with AQ. These data suggest a supra-modal, potentially decision-based, source of this noise.

One possible origin of fluctuations in decision variables could be attentional lapses. To explain the previously reported data (Manning et al., 2015; Manning et al., 2017), we hypothesise that people with higher number of autism traits are more focused and have fewer attentional lapses (i.e., a lower lapse rate), consistent with an earlier report (Bach & Dakin, 2009). This would mean they will have a lower induced noise, which will lead to lower detection thresholds at high external noise levels, but little change in detection thresholds at low external noise levels.

Overall, our discussion suggests that a major difference between people with ASD and typically-developing people may be in their decision process during perceptual-decision making tasks, which can be quantified using induced noise measurement.

### Performance levels

We show no evidence for a uniformly impoverished motion processing in individuals with increased levels of autism traits, which is consistent with the literature. We find that accuracy is generally lower in individuals with higher levels of autism traits, but this was not significant. In the literature, there is evidence for both increased (Foss-Feig et al., 2013; Manning et al., 2015) and decreased (e.g., Koldewyn, Whitney, & Rivera, 2010; Milne et al., 2002; Spencer & O’Brien, 2006) motion perception in ASD (see also Simmons et al., 2009).

There is a stronger case for impoverished motion perception in a particular type of global motion perception, namely biological motion processing. Here too, there is both evidence in favour of (e.g., Blake, Turner, Smoski, Pozdol, & Stone, 2003; Koldewyn et al., 2010; van Boxtel, Dapretto, & Lu, 2016; van Boxtel, Peng, Su, & Lu, 2017), and against (e.g., Cleary, Looney, Brady, & Fitzgerald, 2014; Cusack, Williams, & Neri, 2015; Saygin, Cook, & Blakemore, 2010) a deficit in processing, but a recent meta-analysis found that there was a small decrement in ASD versus typically developing individuals (Van der Hallen, Manning, Evers, & Wagemans, 2018). It may therefore be worthwhile to look at the influence of internal noise on biological motion (as done in van Boxtel & Lu, 2015), while focusing on the link to ASD.

### Motion Pooling

Previous literature has provided both evidence for and against a difference in motion pooling in ASD versus typically developing individuals (Manning et al., 2015; Pellicano et al., 2005). We found no significant correlation between motion pooling across AQ score, and Bayesian statistics suggested that there was moderate evidence for the absence of a correlation in both discrimination tasks. As mentioned above, the increased motion pooling in ASD, reported in previous studies (Manning et al., 2015; Manning et al., 2017), is instead potentially attributable to decreased induced noise (or increased noise exclusion).

### Coarse versus fine discrimination tasks

A reduction of induced noise in individuals with more autism traits, is consistent with recent reports of superior motion perception in ASD (Foss-Feig et al., 2013; Manning et al., 2015). However, several other reports show inferior performance in ASD (e.g., Koldewyn et al., 2010; Milne et al., 2002; Spencer & O’Brien, 2006), especially in motion coherence tasks (Simmons et al., 2009). These differences could be due to the large variability across individuals falling on the autism spectrum, but they could also be due to the different parameters used in the various studies. Indeed, our results support the suggestion that small parameter differences can impact the results, as we found that, while internal noise was higher for individuals with more autism traits in the coarse discrimination task, this was not the case in the fine motion discrimination.

What are the processing differences that could underlie the different findings in our fine and coarse discrimination tasks? Because the stimuli in our two experiments were nearly identical, the difference cannot be attributed to some general motion perception deficit, or other commonly suggested deficits in ASD, such as, a dorsal stream deficit (Grinter, Maybery, & Badcock, 2010; Spencer et al., 2000), a magnocellular dysfunction (Sutherland & Crewther, 2010), or difficulty with complex stimuli (Bertone et al., 2003; Bertone, Mottron, Jelenic, & Faubert, 2005).

Rather our results do align with the more general notions that individuals with ASD are more “detail-focused” (Happé & Frith, 2006; Mottron, Dawson, Soulieres, Hubert, & Burack, 2006), and only show increased internal noise in the coarse discrimination task. An alternative, not mutually exclusive, account is that different ‘decoding rules’, or decision rules, are used for fine and coarse discrimination tasks (Jazayeri & Movshon, 2007), and that they are differentially affected by noise, and autism traits. This suggestion is supported by our comparison of internal noise values between the two tasks. We find that internal noise is larger in the fine discrimination task. Our data suggest that the type of task influences the amount of internal noise, even with near identical stimuli. Although, caution is warranted, as we used different participants in both experiments.

### Conclusions

This research is the first to attempt to relate Autism Spectrum Questionnaire (AQ) scores to estimates of additive and induced noise limits on motion direction discrimination. We show influences of additive and induced noise as well as motion pooling on motion perception. Internal noise is higher in individuals with more autism traits, but the individual noise measures of additive noise, and motion pooling, and induced noise do not correlate with ASD traits. This suggests that a combination of these three (and potentially other) factors, increases internal noise, and that there is no individual type of noise that is solely responsible for the overall increase in internal noise.

We ascribe induced noise to variability in decision-making, and argue that this could provide an alternative explanation of past results indicating superior motion averaging in ASD. The involvement of induced noise in motion perception is very relevant to ASD research, because induced noise affects the perception of supra-threshold stimuli, as opposed to additive noise which mostly affects peri-threshold perception. This implicates internal, and specifically induced, noise as an explanation for (perceptual) atypicallities in ASD, including hyper- and hyposensitivity.

## Acknowledgements

The authors thank Tori Gaunson, and Mary Abad for help with the data acquisition.

## Author contributions

EO and JVB conceived of the experiment, EO and JVB implemented and ran the experiment. EO and JVB performed the data analysis. EO wrote the first draft, JVB wrote the final draft. SD aided in the interpretation of the results, and provided feedback on the final draft.

## Appendix

We performed a simulation study to investigate which approach best estimates the different sources of internal noise and motion pooling. We used 100 double-pass trials (200 trials total), just as for our experimental data. We set additive noise, induced noise and motion pooling to reasonable values. If the final activity (that is signal plus noise) was larger than zero, a “right” response was recorded, otherwise a “left” response was recorded. Accuracy and consistency values were calculated, and then fitted with the same procedures as in the experiment. This resulted in internal noise estimates at each external noise level. Using the following simulations, we aimed to determine which method was best able to extract the original additive noise, induced noise and motion pooling values. The code for these simulations can be found at: https://osf.io/4gdkt/.

The first approach was to fit one function to the whole internal noise versus external noise curve (like our data in Figure 6 and Figure 9). According to the definition of induced noise (Burgess & Colborne, 1988) total internal noise is equal to 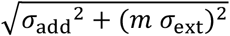. This definition however does not include the possible effect of motion pooling, which we observed in our data, so we used the following function (see equation (*1*)):

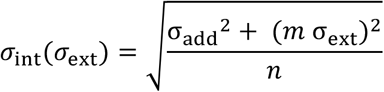

We tested several ways of fitting this function. First, we let all parameters (*σ*_add_, *m*, and *n*) vary freely. Second, we estimated *n* from our data, using equation (2), and estimated only *σ*_add_, and *m*. The third and fourth approach were based on the following approximation: as σ_ext_ becomes large, σ_int_ approaches *m* σ_ext_, which means that *m* = σ_int_/σ_ext_, that is, the σ_int_/σ_ext_ ratio that we measure with the double-pass paradigm. This method ignores the influence of motion pooling, which can be included and would lead to the following estimate: 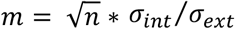. We found, however, that this latter estimate severely overestimates *m*, and we do not discuss it here. The third method derived σint/σ_ext_ by fitting σ_int_/σ_ext_ curves through each of the data points from the largest 5 external noise values, and taking the median value of these fits. The fourth method derived σ_int_/σ_ext_, fitting one σ_int_/σ_ext_ curve (c.f., Figure 4) through the data from the largest 5 external noise values only, and this ratio was taken as the induced noise factor *m*.

Each simulation performed 500 runs, resulting in a distribution of extracted values for additive noise, induced noise, and motion pooling. We then performed different simulations and checked which approach overall appeared to give the best estimates of the inputted additive noise, induced noise, and motion pooling.

Figure 11 shows the results for one of these simulations for the small angle experiment. It shows (as most simulations did) that the best approach was to estimate pooling from the data, and fit additive noise and induced noise (panel b, the WOP fit, that is the fit without motion pooling). The approach in which all parameters were fitted was very dependent on initial conditions (this was because many parameter combinations led to virtually identical fits). The other two approaches overestimated induced noise (and did not provide estimates for additive noise and pooling). The simulations for the large angle data showed similar findings, but also indicate that additive noise could not be estimated very well for this experiment (not shown). We therefore decided not to do statistics on the additive noise measures for the large angle experiment.

**Figure 11.**
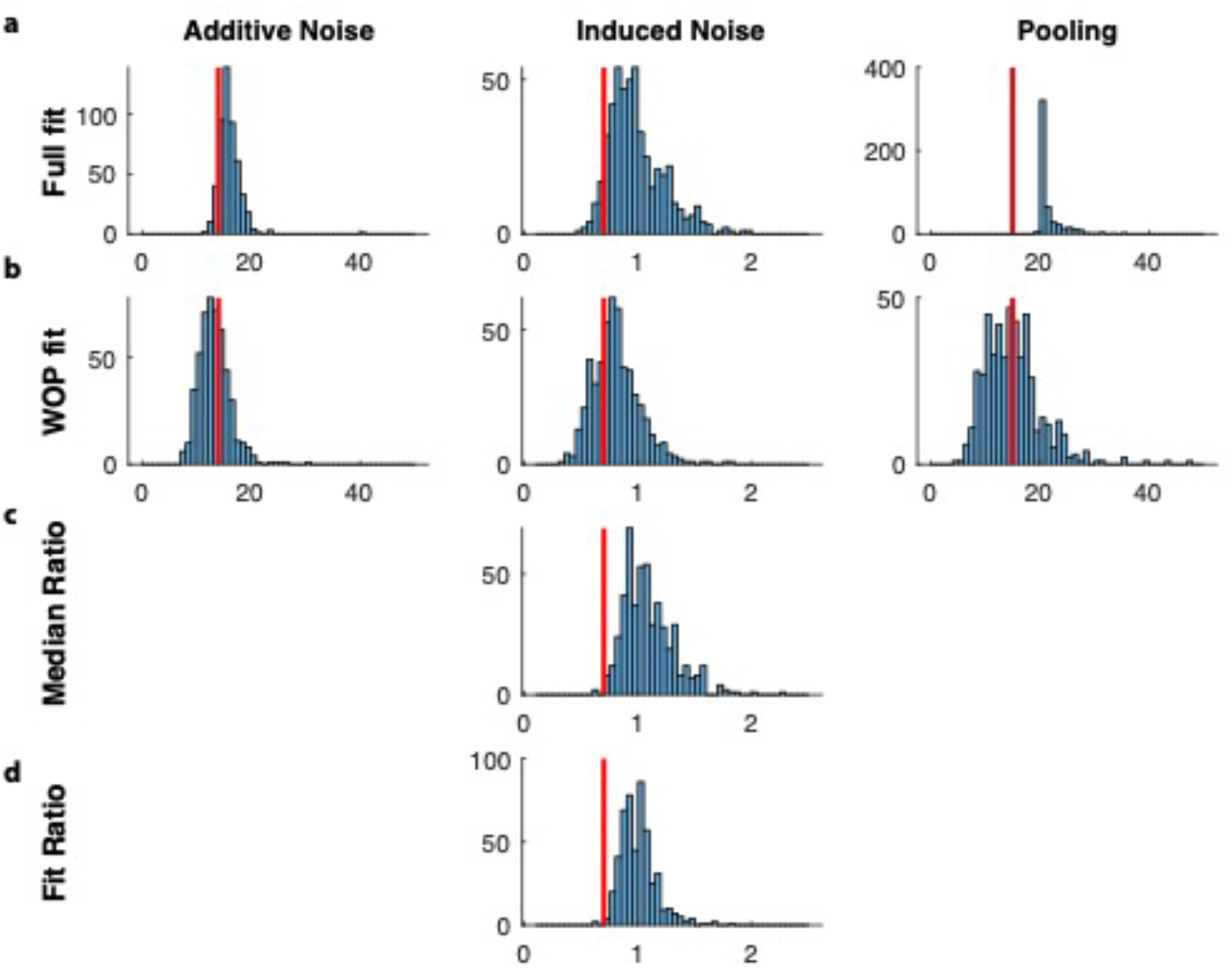
Simulation results for different fitting procedures, for the fine direction discrimination experiment. Results show histograms of estimated model-parameters from 500 simulated runs. The red lines indicate the true values. (a) Full fit in which additive noise, induced noise, and motion pooling were all individually fit. (b) Fit without pooling (WOP) in which additive noise and induced noise were fitted, but the pooling parameter was estimated directly from the data. (c) Estimated induced noise; the median induced noise value for the 5 largest external noise conditions. (d) Estimated induced noise taken from a single fit through the 5 largest external noise conditions.

These simulations also indicated that motion pooling could be estimated from raw data using Equation 2, but only for higher external noise values for the small angle experiment (Figure 12a), and for the lower external noise values for the large angle experiment (Figure 12b).

**Figure 12.**
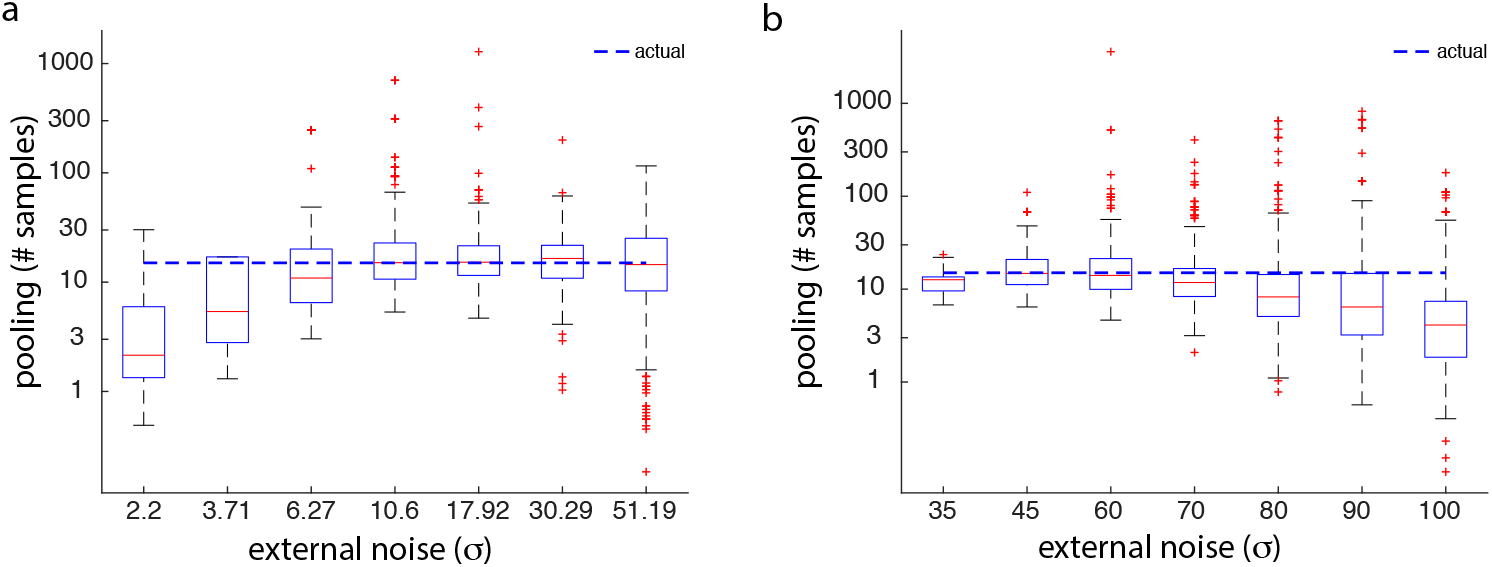
Estimated motion pooling sample-size, based on equation 2 from one of the simulations. (a) Estimates for the fine direction discrimination task is accurate for larger external noise values used in the experiment. (b) Boxplots for the coarse direction discrimination task. These estimates are accurate for lower external noise values used in the experiment. The box of the boxplots contains the 25^th^-75^th^ percentiles, the whiskers show the most extreme values not considered outliers, and the red markers show the outliers.

## Footnotes

1 Induced noise may also be interpreted in terms of “noise exclusion” (i.e., the ability to have external noise not influence internal noise). In this article, we will not distinguish between these different processes. Another type of noise mentioned in the literature is multiplicative noise, which scales both signal and noise. In our paradigm, this behaviour is interpreted as motion pooling.

